# Sweet interference: oral fermentation volatile confounders in exhaled breath revealed by minimal glucose exposure

**DOI:** 10.64898/2026.06.10.731394

**Authors:** Anesu Chawaguta, Daniel Sanders, Veronika Ruzsanyi, Chris A. Mayhew, Lorenzo S. Petralia

## Abstract

Volatile organic compounds (VOCs) in human breath have been explored as non-invasive biomarkers for disease, including respiratory infections and cancer, yet very few breath tests have reached clinical validation and regulatory approval. A major barrier is the difficulty of identifying and controlling confounding factors that affect volatile exhaled breath composition. A critical and overlooked confounder is the oral microbiome, which produces VOCs that can obscure the trace volatiles originating from the lower airways. To investigate this, we conducted an intervention study on sixteen healthy volunteers, using real-time breath analysis, which demonstrates that oral microbiota rapidly alter exhaled VOC profiles following a low-dose (0.5 g) oral glucose administration. Acetoin levels respond promptly to glucose, confirming its oral microbial origin. However, pathogenic bacteria resulting from respiratory infections can also produce acetoin, underscoring the challenge of distinguishing sources of breath VOCs. Similarly, other volatiles, such as acetic acid and ethanol, are also influenced by small glucose doses, complicating their use as biomarkers in non-targeted volatilomic studies. Recognising the metabolic context of each volatile is essential to distinguish infection signals from physiological background. Beyond serving as a cautionary note for exhaled breath research, these results may encourage the oral health and dentistry communities to adopt breathomics analytical tools for rapid chairside diagnostics, transforming respiratory confounders into clinical opportunities for dental care.

## 1 Introduction

Exhaled breath contains a rich-array of volatile organic compounds (VOCs) that reflect human metabolic and microbial processes, making it an attractive non-invasive biomatrix for clinical diagnostics [1, 2, 3]. Despite this promise, clinical translation faces significant challenges. Persistent challenges include a lack of standardisation in breath collection [4, 5] and analytical methodologies, alongside numerous confounding factors (e.g., diet [6, 7, 8, 9], oral-hygiene [10, 11], medication use, and environmental exposure [12]) that may obscure disease-specific signals. Among these confounders, oral microbial contributions to breath VOC profiles are critically underappreciated [13, 14, 15, 16].

Microbial VOCs generated within the oral cavity can profoundly influence exhaled breath composition [17], a phenomenon most clearly illustrated by intra-oral halitosis. Approximately 80-90% of halitosis cases originate from oral bacterial metabolism, primarily driven by anaerobic bacteria residing on the tongue and within dental plaque, which degrade substrates into volatile compounds [18, 19, 20]. Sulphur-containing compounds, such as hydrogen sulphide and methyl mercaptan constitute the dominant malodorous fraction, and are associated with clinical measures including periodontal pocket depth and bleeding [21]. Distinct candidate VOC signatures have also been reported for periodontal pathogens and cariogenic microorganisms using in vitro headspace analysis [22, 17]. Additional VOCs including short-chain organic acids, alcohols, aliphatic polyamines (such as putrescine and cadaverine), and indole further contribute to oral malodour. Several oral anaerobes have been shown to produce hydrogen cyanide (HCN) in vitro [23], while stimulated oral-fluid HCN is significantly correlated with mouth-exhaled HCN and decreases after oral disinfection [24].

When oral microbial activity is elevated, these microbial VOCs can overwhelm breath composition in two distinct ways. First, by elevating the background level of compounds that are also produced systemically, thereby reducing the analytical sensitivity for low-abundance systemic volatiles. A classic example is ammonia, which is generated in the oral cavity via bacterial urease-mediated hydrolysis of salivary urea [25] and is present at substantially higher concentrations in exhaled breath from the mouth than from the nose [14]; its potential clinical utility as a systemic breath biomarker (e.g., for renal or hepatic impairment) is correspondingly complicated by this oral contribution [26], and the real-world diagnostic value of breath ammonia measurements remains actively debated [27]. Second, oral microbial activity may complicate measurements by non-specifically elevating compounds that are also produced by respiratory pathogens (e.g., acetoin [28]), thereby generating false-positive signals and reducing diagnostic specificity.

Compounds such as hydrogen sulphide, 1-propanol, 2-propanol, ethanol, and acetoin are present at significantly higher levels during oral exhalation compared to nasal breathing, demonstrating that the sampling route critically affects detected volatile profiles and that oral-derived compounds can confound biomarkers assumed to originate in the lungs or systemic circulation [16, 29]. Streptococci, among the most prevalent constituents of the oral microbiota, play a central role in this process by mediating in extensive interbacterial and interkingdom interactions that govern biofilm architecture and function. *In vitro* studies show that *Streptococcus mutans* readily ferments glucose to produce ethanol, acetic acid, formate, and acetoin under diverse environmental conditions [30]. Oral bacteria can thus elevate the exhaled breath levels of such VOCs that are often reported as candidate biomarkers for gastrointestinal disorders, liver disease, and respiratory infections [31]. Without explicit consideration of oral microbial sources, associations between these VOCs and systemic disease may be spurious. These considerations underscore the need for improved breath-sampling protocols that mitigate oral microbial confounding effects [16].

Exhaled VOCs can offer valuable insights into host-microbiome-pathogen dynamics during respiratory infections [32, 33]. Given that viruses lack intrinsic metabolic pathways, virus-associated metabolic alterations often arise indirectly, either through host immune responses or through interactions with co-occurring pathogenic bacteria, manifesting as changes in exhaled breath VOC profiles [34, 35, 36].

Among microbially produced VOCs, acetoin (3-hydroxy-2-butanone) has attracted attention as a putative marker of bacterial infection in the respiratory tract. Elevated levels of breath acetoin have been associated with specific pathogenic bacteria [37, 38] and antimicrobial-resistant carbapenemase-producing *Klebsiella* infection [39]. However, while acetoin in breath may signal bacterial infection or superinfection within the airways, its interpretation is complicated by its broad microbial origin. Acetoin is an intermediate in several fermentation pathways [40, 41] and is produced by a wide range of microorganisms. This principle underlies the classical Voges-Proskauer (VP) assay, which distinguishes VP-positive bacteria according to their capacity for acetoin production [42]. Studies have further shown that supplying exogenous sucrose increases pyruvate-derived metabolites, including acetoin [43, 44]. In the oral cavity, increased sugar availability rapidly stimulates acetoin synthesis by resident microbial biofilms [44] and is accompanied by rises in other fermentative VOCs, such as ethanol [45] and acetic acid [46]. Together, these processes highlight how oral microbial activity can substantially influence exhaled breath VOC profiles, complicating the interpretation of breathomic data. To illustrate this complication, consider acetoin, which is a VOC associated with infections in the respiratory tract. In cystic fibrosis, two recent studies ([47, 48]) highlighted changes in acetoin levels in trophic cooperation between *Staphylococcus aureus* (producing acetoin [28]) and *Pseudomonas aeruginosa* (metabolising acetoin). This underscores the complexity of microbial interactions in chronic infections and interspecies cross-feeding mechanisms. Additional studies highlight that infection-associated VOC alterations are not limited to acetoin. Smith et al reported markedly elevated acetic acid concentrations in the breath of cystic fibrosis patients compared with healthy controls, independent of *P. aeruginosa* infection [49, 50]. Similarly, Schnabel et al. identified a panel 12 VOCs capable of discriminating ventilator-associated pneumonia (VAP) with approximately 75% sensitivity and specificity [51]. Within this panel, ethanol was significantly elevated in VAP-positive patients, likely reflecting contributions from both bacterial and host metabolic processes.

These observations exemplify a central challenge in breathomics: many VOCs are produced by both pathogens and commensals, with important consequences for the detection of respiratory infections using exhaled breath-based diagnostics [52, 53]. Not only does the dual origin of these volatiles complicate their biological interpretation, but it also prompts researchers to critically evaluate breath sampling approaches, favouring, for example, nasal exhaled breath over oral samples in order to minimise confounding effects from the oral microbiota [16].

To demonstrate this duality of volatile origins, we report the results of a controlled glucose intervention study conducted with fasting volunteers, designed to assess whether oral microbial activity can confound infection-associated VOC biomarkers in breath-based diagnostics.

## 2 Methods

Sixteen healthy volunteers were recruited for this study. Each participant received an individually sealed, commercially available 2 g glucose candy (”Traubenzucker”, lemon-flavoured), which was quartered into 0.5 g portions. Participants were instructed to chew only a single 0.5 g glucose portion thoroughly to produce a glucose-rich saliva slurry, which was aerated within the oral cavity prior to swallowing. This procedure served as a modified *in vivo* VP test, a well-established microbiology assay used to characterize bacteria by detecting acetoin production following glucose administration [40, 42]. The glucose intervention was designed to probe local fermentative processes within the oral cavity rather than systemic metabolism. Consequently, a dose of 0.5 g was administered, approximately 150 times lower than the standard 75 g bolus used in oral glucose tolerance tests [54, 43, 55], which are intended to assess the systemic glycaemic response. To place this amount in a broader dietary context, 0.5 g of glucose corresponds to roughly 1-2% of the 25-50 g range of daily free sugar intake implied by World Health Organization guidance for adults [56], which is substantially less than the sugar content of a single sugar cube (approximately 4 g) or a few millilitres of a sugar sweetened beverage (which typically contains around 5-10 g of sugar per 100 mL in European surveys). Such sub gram quantities can plausibly be supplied by residual food particles retained in interdental spaces or on mucosal surfaces between meals [57] indicating that the fermentative responses characterised here reflect a physiologically realistic oral sugar exposure rather than a supraphysiological challenge.

To minimize potential confounders and ensure data quality, participants were excluded if they had consumed alcohol within the previous 24 hours, recently used antibiotics or antiseptic mouthwash, exhibited signs of acute infection, smoked or vaped within 30 minutes before sampling or consumed food or sweetened beverages within two hours prior to breath sampling.

Exhaled breath was spectrometrically analysed in real-time using a Proton Transfer Reaction-Time-of-Flight-Mass Spectrometeter (PTR-ToF-MS) from Ionicon Analytik GmbH, specifically a PTR 6000 X2, using H_3_O^+^ as the reagent ions [58]. PTR-ToF-MS is a well-established technique for real-time VOC analysis, and its analytical principles are well documented in the literature [59, 60] ; therefore, only the operational parameters used in this study are described here. The inlet-system and the drift tube were maintained at a temperature of 100 °C; the drift tube was operated at a pressure of 2.6 mbar. All measurements were conducted in DC-mode (no RF ion funnel) at a reduced field strength of 120 Td. The inlet system was interfaced to a buffered end-tidal (BET) sampling device [61], which allowed the end-tidal portion of each exhalation to be easily sampled. The BET was operated at 70 °C in order to prevent condensation of humid breath and to minimise adsorption of volatile compounds on the internal sampling surfaces.

Participants were seated comfortably near the instrument for 5–10 minutes before sampling to achieve a stable physiological baseline and minimise the risk of hyperventilation. They were then instructed to exhale approximately once per minute through a disposable, one-way mouthpiece connected to the BET device. After three exhalations, the oral glucose was administered, after which participants continued exhaling into the BET at one-minute intervals for a further 25-minute monitoring period (see **Figure 1** for a schematic overview of the protocol) .

**Figure 1.**
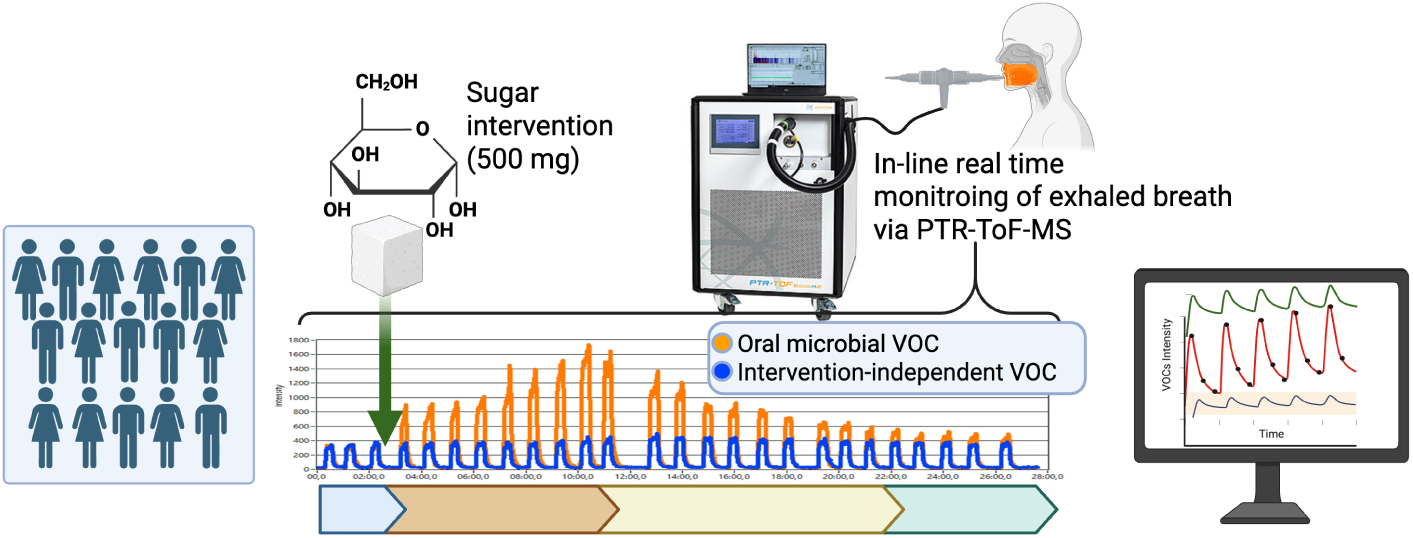
Schematic overview of the study protocol. Each volunteer’s oral end-tidal breath was continuously monitored using PTR-ToF-MS and the BET sampling interface. After the first three measured exhalations, participants were administered 0.5 g of glucose, after which breath measurements continued at regular intervals for approximately 25 minutes. The coloured bar at the bottom represents the pre intervention phase, the initial post intervention rise, the peak response, the subsequent decay phase and the final post intervention plateau observed within the experimental time window. The distinction between “oral microbial” and “intervention independent” VOC classes is shown schematically and draws on previous comparative oral and nasal breath sampling studies [16]. Image created in BioRender.

### Data Analysis

PTR-ToF-MS spectral data, including peak positions (*m/z* values) and signal intensities, were analysed using the manufacturer’s software PTR-MS Viewer (v3.4.5; Ionicon Analytik GmbH). Given that this study was strictly hypothesis driven, no untargeted feature set was generated, and our inferential statistical analysis focused on an a priori selected panel of seven target volatile organic compounds (ethanol, acetic acid, acetoin, 2,3-butanediol, limonene, acetone and isoprene). These variables were chosen based on their established relevance to oral fermentation chemistry and systemic breath physiology, ensuring that the subsequent statistical tests remained mechanistically interpretable (see also **Table S1** in the Supplementary Material for more details). Notably, the presence of limonene associated with the citrusy flavouring of the candy provided a distinct exogenous breath signal, allowing for a clear identification of the intervention time point. The volatile compounds monitored in this study were identified in preparatory work using gas chromatography-ion mobility spectrometry (GC-IMS); relevant analytical information is provided in the Supplementary Material (see Table S2 and Figures S1, and Figure S2). PTR-ToF-MS was subsequently adopted for its high time resolution, enabling real time monitoring of the intervention response.

For each participant, PTR-ToF-MS scans were processed at five important time points, namely *t_before_*, *t*_0_, *t_p_*, *t*_8_, and *t*_end_, which correspond to the exhalation prior to the glucose intervention, the breath sampled immediately after glucose ingestion, the breath when acetoin peaked, the breath collected 8 minutes after glucose administration, and the final recorded exhalation (as shown in **Figure 2**).

**Figure 2.**
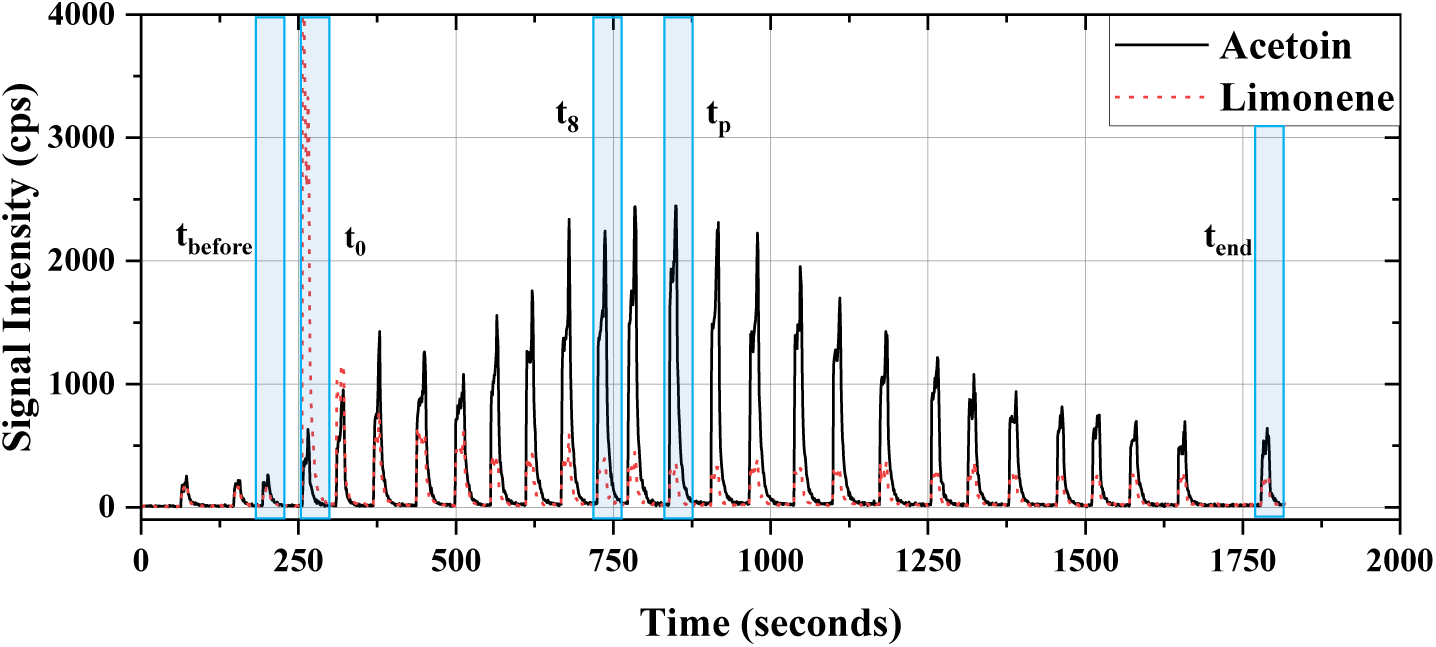
Time-resolved breath VOC response to glucose administration in a representative participant. End-tidal concentrations of acetoin (black line) and limonene (red dotted line) measured by PTR-ToF-MS continuously over approximately 30 minutes. Acetoin exhibits a rapid rise within minutes following glucose administration, followed by a gradual decline, consistent with a fast and transient oral microbial fermentation response.. In contrast, limonene shows an immediate post-intervention peak due to the flavour additives present in the glucose sweet. Five characteristic time points are selected for the inter-subject analysis, namely *t_before_*, *t*_0_, *t_p_*, *t*_8_, and *t_end_*, as defined in the text.

Subsequent data parsing and statistical analyses were performed in Python [62, 63], using a workflow combining multivariate pattern exploration with repeated-measures inference. Principal component analysis (PCA) was deployed as an exploratory tool to visually inspect the multivariate temporal structure of the dataset prior to applying robust multiple comparison statistical tests. Prior to the statistical analysis, the VOC signals were log-transformed to stabilize variance and reduce the right-skewness typical of mass-spectrometric signals. Multivariate structure was investigated using PCA with standard-scaled data. PCA score bi-plots were complemented by confidence ellipses representing the covariance structure of each time point, enabling visualization of within-group variability and separation across the intervention. Loading vectors were examined to identify the VOCs contributing most strongly to temporal differentiation.

For each VOC, where assumptions were met, repeated-measures ANOVA was applied; otherwise, non-parametric Friedman test was used. Effect sizes were quantified using partial eta-squared (*η*^2^) for parametric tests and Kendall’s *W* for Friedman tests. Omnibus significance was followed by pairwise Wilcoxon signed-rank tests between time points. Multiple-comparison corrections were applied using the Holm–Bonferroni procedure, which controls the family-wise error rate (FWER). Rank-biserial correlation was reported as the effect size for the various pairwise contrasts.

## 3 Results

Real-time monitoring of oral end-tidal exhaled breath from sixteen healthy adults revealed dynamic changes in specific VOC concentrations following the chewing and then ingestion of 0.5 g of glucose. These alterations were monitored in real-time over an approximately 25-minute period post chewing the glucose.

Identifying statistically significant changes in individual VOCs proved non-trivial. The detection of subtle but biologically meaningful variations within highly variable datasets required careful statistical handling, and the minimisation of major confounding influences. This underscores some of the inherent challenges associated with exhaled breath volatile biomarker discovery in the presence of microbial variability.

Inter-subject variability in terms of the temporal evolution and levels of the specific VOC concentrations was observed (as also shown in **Figure 3**). This means that the use of fixed VOC concentration cut-off values is impractical. Moreover, a subset of volunteers did not exhibit a pronounced post-glucose increase in acetoin or ethanol despite identical experimental protocols. These inter-individual differences could reflect differences in oral microbial composition and metabolism, including differences in homo-/heterolactic fermentation, local pH conditions [13], and the availability of heme [64] or molecular oxygen (see Discussion section).

**Figure 3.**
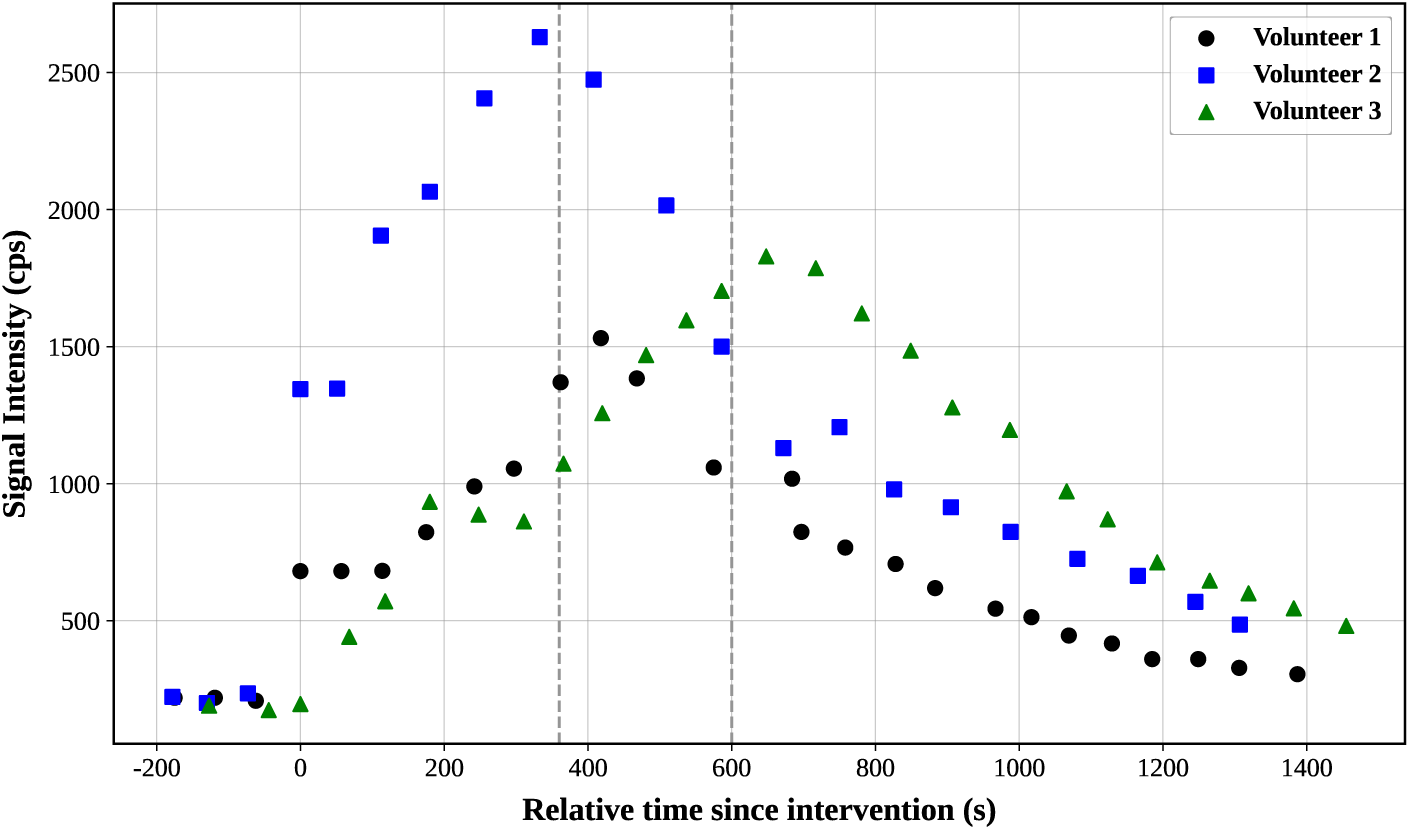
Representative acetoin time-courses from three participants following the glucose intervention. Volunteer 1 featured an acetoin peak close to the average time *t*_8_; whereas volunteer 2 and volunteer 3 showed the earliest and the latest acetoin peaks, respectively. Owing to inter-individual variability in oral-microbial metabolism and breath-biome dynamics, the time point at which acetoin reached its maximum concentrations, *tp*, was defined individually for each participant. Across participants, peak acetoin production occurred approximately between 6 and 10 minutes (vertical dashed lines) after administering glucose.

**Figure 3** shows the temporal evolution of acetoin concentrations curves for three representative participants. Owing to inter-subject variability arising from differences in host metabolism and oral microbiome composition, and salivary flow (with inevitable variations in chewing/swallowing), a fixed post-intervention time as to when the acetoin reached a maximum value does not occur. Instead, we define individual time points, *t_p_*, at the moment when the acetoin signal in the exhaled breath of a volunteer reached its maximum. This approach enabled the capture of the peak metabolic response to glucose exposure and facilitated pre- and post-intervention comparisons independently of subject-specific response kinetics. The use of end tidal breath rather than salivary headspace or low flow oral breath sampling further indicates that the observed differences were not simply generated by sampling an undiluted oral headspace. Exhaled acetone and isoprene levels remained stable throughout the measurement period, indicating that (as expected) true end-tidal concentrations of these compounds were not affected by the glucose intervention.

PCA, employed to investigate the multivariate structure, revealed that, in addition to limonene originating from the glucose candies, the loading vectors most strongly driving the separation between pre- and post-intervention breath samples (both at *t*_p_ and at the *t*_8_ time points) are those of acetoin, ethanol, and acetic acid (see **Figure 4**). In contrast, the loading vectors for isoprene, acetone, and 2,3-butanediol are oriented approximately orthogonal to the microbiome associaied VOCs, indicating that they contribute to variance largely independent of the glucose intervention.

**Figure 4.**
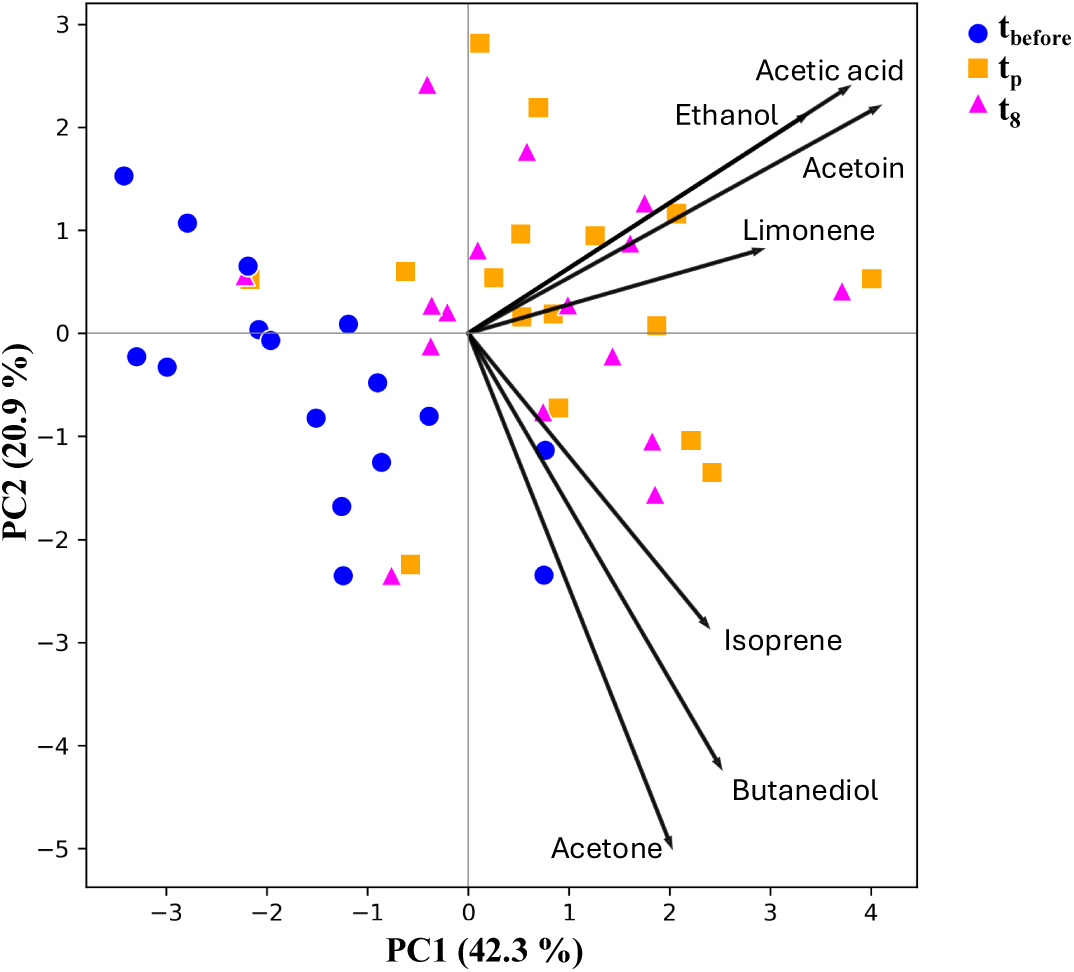
PCA biplot of breath VOC profiles before and after the glucose intervention. The two principal components are shown on the x- and y-axis, with the percentage of explained variance reported in brackets. In addition to limonene (an exogenous VOC derived from the flavouring agent in the glucose candies), the separation between pre- and post-glucose intervention clusters is primarily driven by acetoin, acetic acid, and ethanol. The loading vectors for these three VOCs are approximately orthogonal to those of isoprene, acetone, and butanediol.

On this basis, we selected acetoin, ethanol, and acetic acid as the three VOCs most strongly driving the group separation and repeated the PCA selecting only this reduced set of features and adding the *t_end_* data (see **Figure 5**). Confidence ellipses were overlaid on the score plot to summarize the multivariate dispersion within each time point and to emphasize the between-time-point separation. The minimal overlap between ellipses for the pre- and post-intervention samples indicates that the glucose intervention induced a consistent shift in the overall VOC profile across participants. The ethanol loading vector predominantly drove the separation of the tend cluster from the earlier time points. This observation identifies ethanol as the principal contributor to the later-phase temporal evolution of the VOC profile following glucose administration.

**Figure 5.**
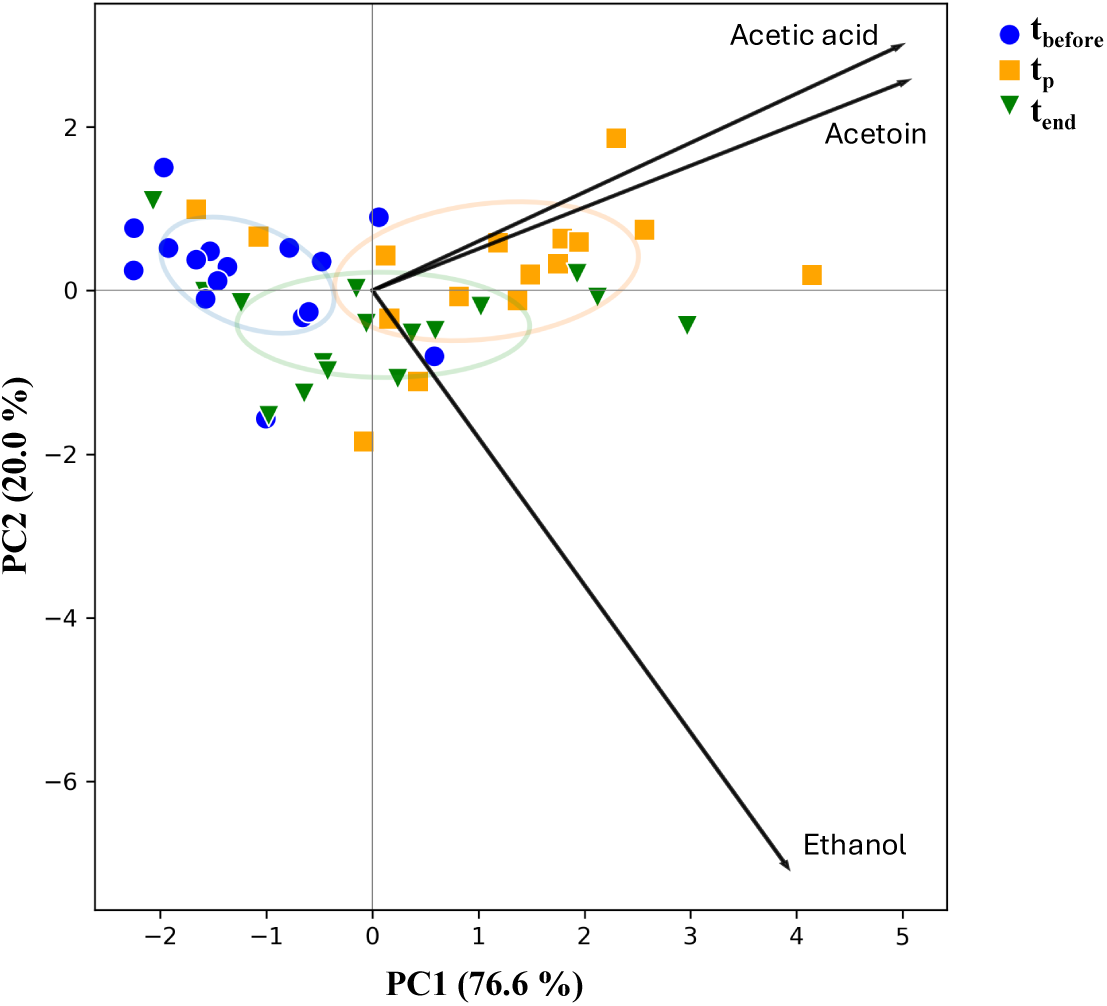
PCA biplot of the *t*_pre_, *t*_p_, and *t*_end_ samples using the feature-selected VOCs (acetoin, acetic acid, and ethanol) based on the previous PCA (see Figure 4). The graph shows improved percentage of explained variance and separation between the pre- and post-intervention time points with negligible overlap of the *t*pre and *t*p confidence ellipses. Notably the *t*_end_ cluster (featuring the last time point samples) is separated primarily along the ethanol loading vector.

The temporal distributions of the measured values across time points demonstrates that, unlike acetoin and acetic acid, ethanol exhibits a post-intervention increase that does not peak (see **Figure 6**). Instead, ethanol concentrations continue to rise, or at least remain elevated, indicating a distinct temporal response compared with the other two glucose-responsive VOCs. Figure 6 also provides values of the isoprene levels during the measurements, which are clearly not affected by the glucose intervention, as expected because it being a purely systemic VOC.

**Figure 6.**
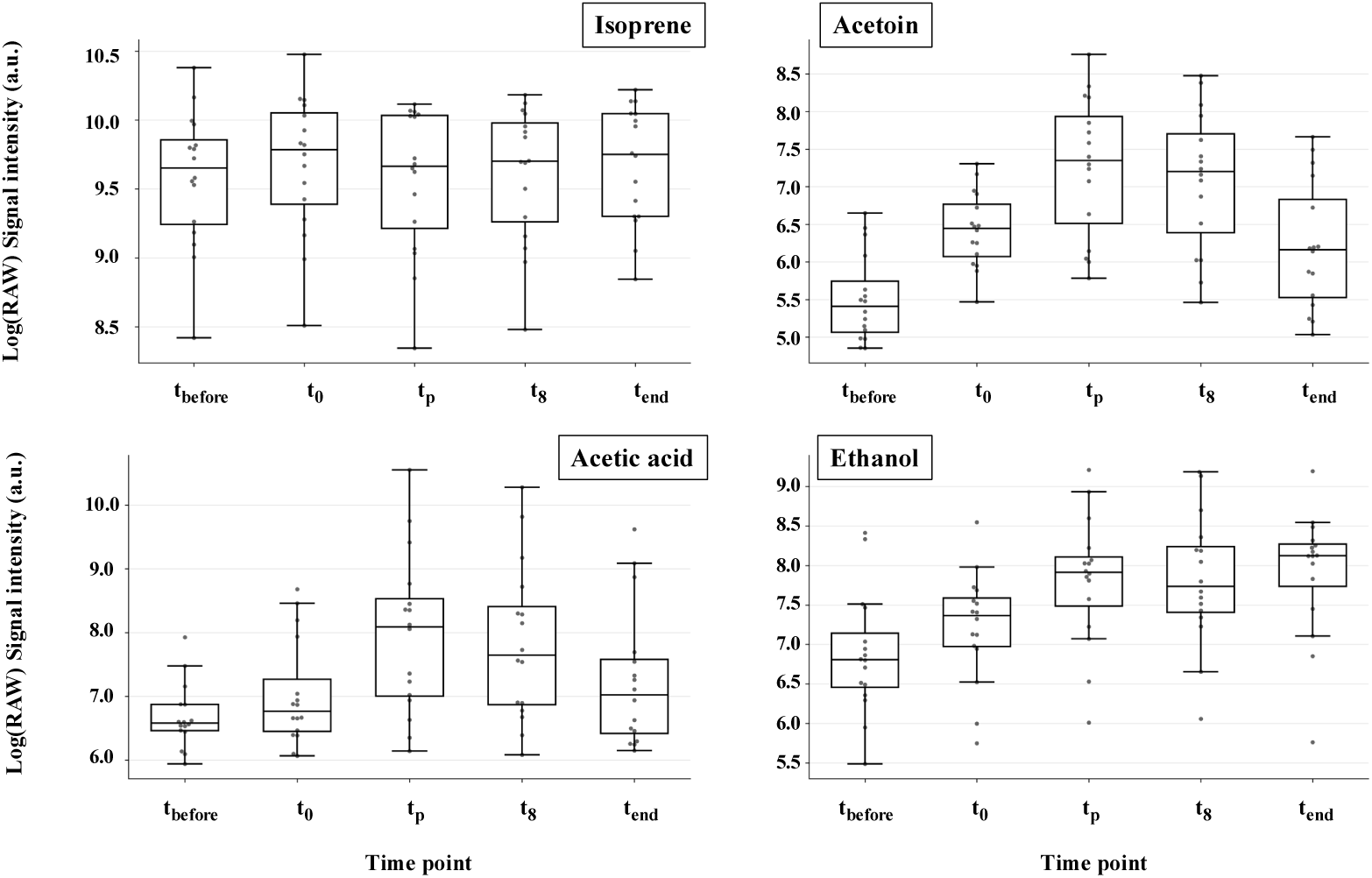
Boxplots of end-tidal VOC signal intensities across the time course of the measurements. Log-transformed signal intensities of isoprene (top left), acetoin (top right), acetic acid (bottom left), and ethanol (bottom right) are shown across the five defined time points for all volunteers. Isoprene, a systemic VOC, remains unaffected by the glucose intervention. In constrast, acetoin and acetic acid exhibit rapid post-intervention increase followed by a subsequent decline, whereas ethanol displays a delayed and more sustained rise. Despite considerable inter-individual variability in absolute signal magnitude, the temporal pattern is reasonably consistent across participants, with acetoin reaching a maximum between 6-10 minutes post glucose administration.

Further robust repeated-measures statistical comparison and post-hoc analysis revealed statistically significant differences between time points (see **Table 1**) regardless of any subject-specific factor (e.g., oral microbiome, oral biofilm, and pH).

**Table 1.** Summary of temporal changes in key VOCs following the glucose intervention. Arrows indicate the direction of change from baseline (in the first two columns), and from *tp* (in the last column). Statistical significance based on Holm–Bonferroni correction (FWER) is denoted as n.s. (*p >*0.05), * (*p <*0.05), ** (*p <*0.01), and *** (*p <*0.001). Effect sizes are reported as rank–biserial correlation coefficients (*r*).

| VOC | Baseline → Peak | Baseline → End | Peak → End |
| --- | --- | --- | --- |
| Isoprene | n.s. | n.s. | n.s. |
| Acetone | n.s. | n.s. | n.s. |
| Butanediol | n.s. | n.s. | n.s. |
| Acetoin | ↑*** ($r = -1.00$) | ↑** ($r = -0.88$) | ↓*** ($r = 1.00$) |
| Acetic acid | ↑*** ($r = -0.97$) | ↑* ($r = -0.79$) | ↓*** ($r = 0.98$) |
| Ethanol | ↑*** ($r = -0.94$) | ↑*** ($r = -0.94$) | n.s. |

Acetoin and acetic acid exhibited the strongest and most reproducible temporal responses to the glucose intervention. Friedman tests showed a highly significant effect of time for both compounds; acetoin (*p* = 1.63 *·* 10*^−^*^10^, Kendall’s *W* = 0.81) and acetic acid (*p* = 3.70 *·* 10*^−^*^8^, *W* = 0.63). Post-hoc comparisons revealed that, for both metabolites, concentrations at *t*_0_, *t_p_*, and *t_end_* were significantly elevated relative to baseline, even after Holm–Bonferroni correction, with very large rank-biserial effect sizes (*|r|* = 0.79 – 1.00). Although the differences between baseline and *t_end_*were smaller than those observed at earlier post-intervention time points, they remained statistically significant, indicating that concentrations had only partially returned toward pre-intervention levels by the end of the monitoring period.

Ethanol showed a significant overall temporal effect (Friedman’s *p* = 1.84 *·* 10*^−^*^7^, Kendall’s *W* = 0.58), but with pronounced inter-individual variability. Post-hoc analyses revealed significant increases in ethanol levels from baseline to *t_p_*, *t*_8_, and *t_end_* (Holm-corrected *p <* 0.001; *|r|* = 0.81–0.94). Ethanol at *t_p_* was significantly higher than at *t*_0_, but did not differ significantly from those observed at *t*_8_ or *t_end_*. This temporal pattern contrasts with the rapid responses observed for acetoin and acetic acid are is consistent with ethanol’s delayed and more sustained production following glucose intervention (see **Discussion**).

By contrast, acetone and isoprene showed no significant temporal variation (all Friedman test *p >* 0.5), consistent with their expected stability as systemic VOC biomarkers. Likewise, 2,3-butanediol exhibited only weak, uncorrected pairwise differences without surviving multiple-comparison correction, indicating a limited and non-robust response to the glucose intervention

## 4 Discussion

PCA consistently highlighted a core subset of VOCs that drove the temporal separation of samples, underscoring the robustness and reproducibility of the observed response pattern. Across participants, a shared directional increase in acetoin, acetic acid, and ethanol was evident following the glucose intervention, despite substantial inter-individual differences in response magnitude. This variability likely reflects the interplay of multiple biochemical and ecological factors within the oral cavity, including differences in microbial community composition, acid tolerance, salivary flow rates, and pH dynamics [13]. Collectively, these factors shape the oral fermentation landscape and modulate the extent to which microbial metabolites are released into exhaled breath.

The repertoire of fermentative pathways available to resident microbes [65], ranging from homolactic conversion of glucose to lactate, to heterolactic production of ethanol and acetate, to the acetoin pathway, means that even identical glucose challenges can yield markedly different VOC temporal profiles depending on the oral pH and which enzymes predominate. For instance, individuals whose plaque microbiomes are enriched in homolactic fermenters may exhibit pronounced lactate accumulation with comparatively modest increases in ethanol or acetoin, whereas those harbouring higher proportions of mixed-acid fermenters are likely to generate elevated levels of ethanol and acetoin under the same conditions [66, 67].

Glucose catabolism proceeds via the Embden–Meyerhof–Parnas pathway (EMP), yielding pyruvate together with a net gain of two ATP molecules and the reduction of two molecules of nicotinamide adenine dinucleotide (NAD^+^) to its reduced form, NADH, per molecule of glucose [68]. Under fermentative conditions, sustaining this flux requires continuous re-oxidation of NADH to NAD^+^ through downstream pathways that reduce organic intermediates derived from pyruvate.

In a range of acetoin-producing bacteria, part of the pyruvate pool is diverted from homolactic or mixed-acid fermentation into the acetoin/2,3-butanediol branch, generating more neutral end-products and thereby moderating net acidification [40, 41]. In this pathway, two molecules of pyruvate are condensed to *α*-acetolactate by acetolactate synthase, which is then followed by decarboxylation to acetoin catalysed by *α*-acetolactate decarboxylase. Under appropriate redox and regulatory conditions, acetoin can subsequently be reduced to 2,3-butanediol by butanediol dehydrogenase (BDH), oxidizing NADH to NAD^+^ and thereby contributing to redox balance.

In our oral glucose investigation, we observed a clear post-stimulus rise in exhaled acetoin, whereas exhaled 2,3-butanediol remained essentially unchanged, despite the known biochemical linkage of these metabolites at the pyruvate node. This pattern is compatible with 2,3-butanediol acting as a more downstream product that requires specific enzyme expression and redox states that may not be substantially engaged during the short-term perturbation examined here. Moreover, 2,3-butanediol is markedly more hydrophilic and less volatile than acetoin [69]; consequently, any 2,3-butanediol formed is likely to be retained within oral aqueous compartments and thus only to contributes weakly, if at all, to the exhaled gas-phase signal under our experimental conditions.

We further observed that the peak concentration level for acetoin was typically reached between 6 and 10 minutes after administering glucose, whereas ethanol exhibited a delayed rise, typically reaching the maximal levels toward the end of the measurement window and, in contrast to acetoin and acetic acid, showed no significant decline thereafter (see **Table 1**). These differing dynamics likely reflect the complex biochemical pathways (see **Table 2**) and the interplay of pH variations within oral biofilms.

**Table 2.**
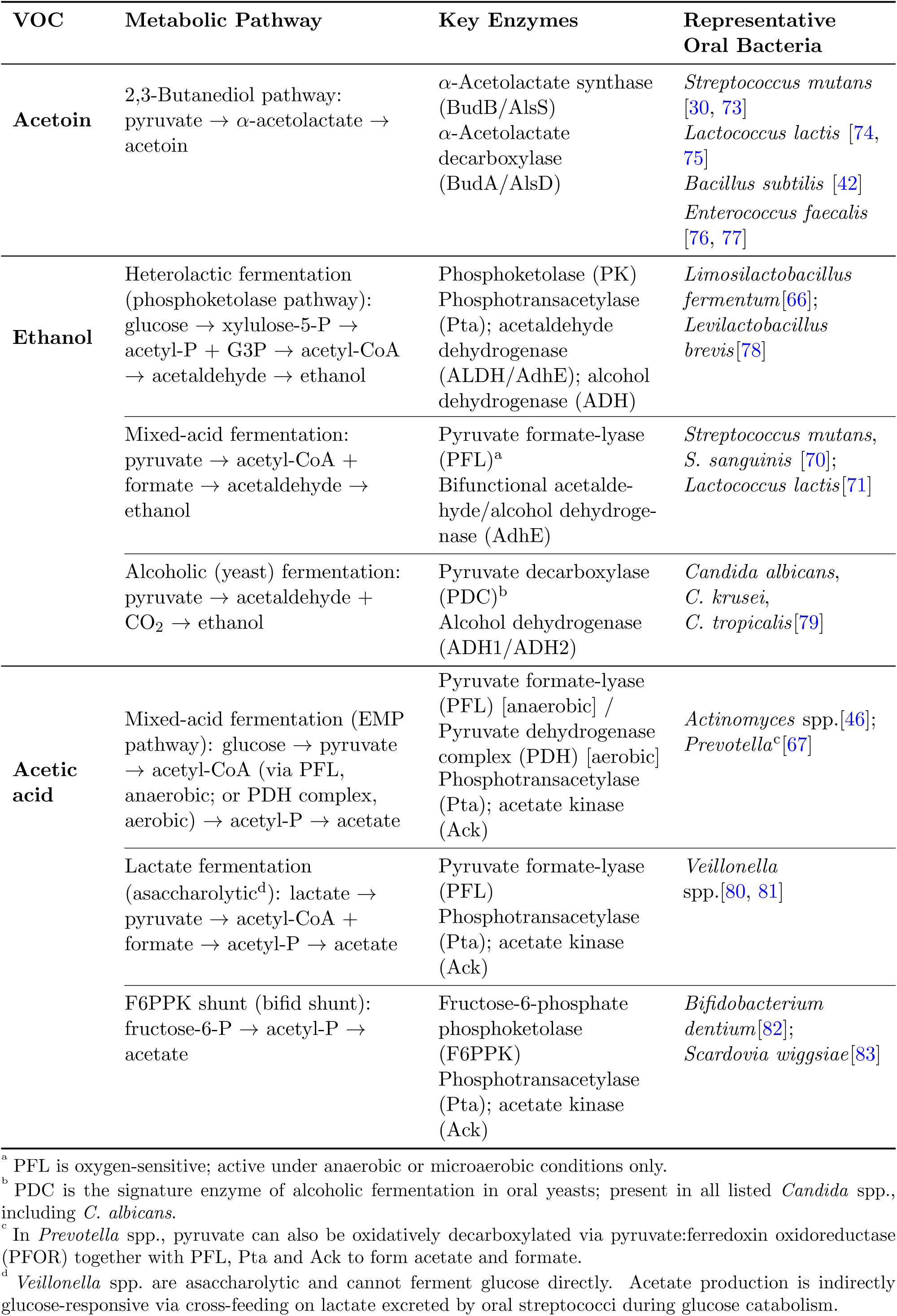
Selected pathways and representative oral microbes linked to glucose-responsive VOC production in the oral cavity.

Following glycolysis, pyruvate can be directed into ethanol formation as part of redox balancing. In many microorganisms, pyruvate is first decarboxylated to acetaldehyde with concomitant release of CO_2_, either by pyruvate decarboxylase or by pyruvate-decarboxylating enzyme systems such as the pyruvate dehydrogenase or pyruvate-formate lyase routes when coupled to acetaldehyde dehydrogenase [66, 70, 71]. Acetaldehyde is subsequently reduced to ethanol by alcohol dehydrogenase, a terminal step that oxidizes NADH to NAD^+^ and thereby sustains glycolytic flux under fermentative conditions. This provides a plausible explanation for why the exhaled ethanol concentrations did not decrease towards pre-glucose baseline values within the observation window.

Notably, a pronounced oral contribution to exhaled ethanol following sugar exposure has previously been reported by Španěl et al, who used SIFT-MS to monitor exhaled breath after mouth rinses with highly concentrated sucrose solutions, at levels exceeding those typically encountered in food and drinks [45]. Taken together, the distinct temporal profiles of acetic acid, acetoin, and ethanol illustrate how fermentative fluxes downstream of pyruvate are dynamically redistributed in response to changing environmental conditions within the oral cavity. These shifts are consequently reflected in the evolving profile of exhaled VOCs.

Recent work by Boisen *et al* provides additional support for the interpretation that glucose-responsive VOC profiles reflect phenotypic differences within the oral microbiome [72]. In their study of caries-active and caries-free children, dental plaque from caries-active subjects exhibited significantly higher acid tolerance and distinct shifts in glucose-derived metabolic products, including altered lactate-to-acetate and lactate-to-ethanol ratios. These metabolic differences were accompanied by enrichment of acidogenic genera such as *Streptococcus* (*S. mutans*, *S. sanguinis*, *S. salivarius*), *Prevotella*, and *Veillonella*. Collectively, these observations demonstrate that oral biofilms vary markedly in both composition and metabolic behaviour, particularly under a glucose challenge. Such variability provides a plausible explanation for the inter-individual variations observed in our real-time VOC responses. They also highlight that plaque phenotype, including acid tolerance and fermentative output, may serve as a biologically meaningful indicator of oral health status, reinforcing the potential for breath-based fermentation challenges as rapid, non-invasive tools for caries-risk assessment.

Compounding this metabolic diversity are microenvironmental parameters, most notably pH [13] and heme availability [64], that modulate enzyme activity and pathway selection. In low-pH niches, bacteria tend to upregulate neutral end-product pathways, such as acetoin synthesis, to avoid intracellular acid stress, whereas more strongly buffered environments favour continued acidogenic fermentation. Moreover, the presence of heme in the oral cavity, particularly under aerobic conditions, promotes a respiratory shift in bacteria such as *L. lactis*, leading to increased acetoin production and improved pH regulation. Such adaptations can influence oral microbial ecology, especially during conditions associated with increased heme availability, including gum disease or oral injury [84]. Taken together, these host-microbiome interactions, shaped by diet, oral hygiene, saliva flow, and individual microbial composition, underlie the broad range of breath VOC responses observed in this study and highlight the critical need to account for these confounders in breath-based diagnostics.

However, a detailed exploration of these factors lies beyond the scope of this present work, which instead aims to bridge research on exhaled volatiles with the oral health and dental science communities. Understanding how shared microbial metabolites such as acetoin, ethanol, and acetic acid arise from both oral commensals and respiratory pathogens strengthens the case for cross-disciplinary approaches in breathomics. To date, the relevance of oral biofilm metabolism and oral microbiota-derived volatiles for oral health and dental science has received only sporadic attention [85]. Oral health status may therefore act as either a confounder in breath-based diagnostics for respiratory disease or, conversely, as an informative diagnostic signal for oral disease itself, underscoring the importance of integrating dental and medical perspectives when interpreting exhaled volatile profiles.

An important physiological note is that glucose is absorbed predominantly in the proximal small intestine, whereas near complete emptying of clear liquids generally requires approximately 30-60 minutes [86, 87]. Colonic fermentation is therefore an implausible explanation for the 1-2 minute onset and 6-10 minute peak observed for acetoin and acetic acid. Moreover, passage of gas from the stomach or more distal gastrointestinal tract to the mouth would also require transient lower oesophageal sphincter relaxation associated with belching [88], and no belching events were observed during sampling.

Another key physiological baseline consideration is that glucose is constitutively present in human saliva [89], even under fasting conditions, sustaining low-level oral microbial fermentation. Whether exhaled acetoin concentrations correlate measurably with blood glucose levels via this salivary glucose route remains an untested but biochemically plausible hypothesis, and one with potential implications for non-invasive metabolic monitoring that warrants direct experimental investigation. Moreover, the observation that even a modest intake of glucose into the mouth, mimicking, for example, residual food entrapped in dental biofilms elicited highly variable exhaled levels of acetoin, ethanol, and acetic acid underscores a twofold implication for breathomics research. First, it serves as a cautionary example that trace amounts of a fermentable substrate in the oral cavity can profoundly skew VOC signatures that are often interpreted as reflecting systemic conditions or lower-airway processes, such as respiratory infections. Without rigorous standardisation of protocols [90], such as controlled fasting intervals, oral rinses procedures, or the selective sampling of end-tidal nasal breath, oral microbial metabolism is likely to remain a major confounder in efforts to assign unambiguously volatile biomarkers to pulmonary pathogens. Nasal breath sampling constitutes a viable approach to mitigate potential confounders, as noted sporadically in the literature [16, 14, 91], but it remains largely overlooked by the broader breath research community.

In contrast, these very confounding influences point to an untapped opportunity in oral health. Oral microbial metabolites contribute to the maintenance of oral homeostasis and are also implicated in oral dysbiosis and disease [65, 92]. Because oral biofilms respond dynamically to fermentable carbohydrate and pH, changes in exhaled VOC profiles following a modest glucose load may provide functional indicators of plaque physiology and oral health status [46, 93, 72]. Tailored breath-based tests (e.g., incorporating bespoke intervention and static sampling) could therefore be developed as non-invasive tools to monitor cariogenic plaque dynamics, dental root canal infections, oral thrush, halitosis, gingivitis, or periodontal disease status [94]. In practical terms, the protocol used here can be regarded as an in vivo analogue of the classical VP test [42], in which a defined substrate is supplied and a neutral fermentation product such as acetoin is detected. Framed in this way, glucose need not be the only stimulus; different substrates would be expected to report on different microbial capabilities, analogous to the classical IMViC panel of biochemical tests (indole, methyl red, VP and citrate), which is used to differentiate bacteria according to distinct metabolic properties [95]. A small panel of such substrate challenges could therefore yield a broader functional readout of oral microbial activity than any single compound.

Volatile metabolites, such as acetoin, may arise from diverse sources, including pathogens, oral commensals, diet, or pH-driven fermentation shifts. Without strict control of sampling routes, oral health, exposure, and dietary factors, breathomics studies (including increasingly common machine learning-based analyses of untargeted volatilomic data) risk misattributing biomarker origins and overstating predictive performance. Researchers must therefore pair advanced analytical tools with rigorous study design, robust sampling strategies, such as end-tidal nasal sampling [16, 90], and mechanistic knowledge of microbial pathways, so that volatile signatures can be safely attributed to disease processes.

## 5 Conclusion

We monitored end-tidal oral breath from sixteen healthy volunteers using PTR-ToF-MS in a controlled intervention study involving the administration of approximately 0.5 g of glucose. Exhaled VOC intensities were tracked over a 30 minute period to characterise the dynamics of glucose-induced oral microbial changes. The measurements revealed a rapid onset of oral fermentation following glucose exposure, while systemic VOCs remained unchanged throughout the experiment. Significant postintervention variations of ethanol, acetic acid, and acetoin were observed in exhaled breath, consistent with rapid microbial fermentation of the small glucose stimulus in the oral cavity. These compounds have also been reported as potential biomarkers of non-oral diseases. Elevated exhaled ethanol and short-chain fatty acids have been linked to gastrointestinal dysbiosis and liver dysfunction, while acetoin has been noted in the breath of patients with respiratory infections. In our intervention study, the pronounced increase in acetoin is instead due to microbial activity within the oral cavity, rather than to systemic metabolic processes.

Although detailed inter-individual differences in oral microbiome composition and metabolic capacity were beyond the scope of this observational study, these findings further substantiate the central role of oral microbial fermentation by showing the marked increases in selected exhaled VOCs following a minute glucose intervention. Notably, even under fasting conditions, small amounts of fermentable carbohydrate may persist within interdental spaces or mucosal niches, providing a readily accessible glucose reservoir capable of sustaining rapid and vigorous microbial fermentation uopn minimal perturbation.

Despite this inter-subject variability, the overall findings highlight a critical consideration for breath-based diagnostics: without stringent sampling strategies and careful control of oral contributions, transient oral microbial fermentation can elevate certain VOCs. Such artefactual increases risk masking or mimicking endogenous, low-abundance biomarkers of systemic origin, thereby compromising the specificity and reliability of breath analysis in clinical contexts.

Breath tests have considerable potential as rapid screening tools for respiratory infections by detecting disease-associated changes in the volatile profile arising from altered metabolism accompanying viral or bacterial infections [52]. However, confounding contributions from oral microbial activity can jeopardise diagnostic specificity and lead to false positive findings. Acetoin exemplifies this challenge: while it is commonly detected in orally exhaled breath as a consequence of glucose-stimulated fermentation by the oral microbiota, its presence in nasally collected breath could serve as a meaningful biomarker of bacterial infection, or bacterial superinfection.

Beyond its implications for standardizing systemic breath analysis, this work also highlights the potential for developing specialized oral-health diagnostic breath tests. Both increased acid tolerance and characteristic microbial consortia have been linked to caries progression, and plaque acidogenic has been proposed as a biomarker for caries risk [72]; moreover, recent work on a methionine challenge test revealed a promising breath-based approach to detect the presence of periodontitis dysbiosis [96]. These findings point toward the future development of targeted breath diagnostics capable of support dental care through rapid, non-invasive assessment of oral microbial activity. To take this work further, targeted breath diagnostic protocols need to be developed for the dental medicine and oral health research communities that exploit oral microbial volatile production as functional markers of dysbiosis, caries susceptibility, gingivitis, and periodontal disease status.

## Acknowledgements

We acknowledge the EU HORIZON Innovation Actions HORIZON-CL3-202-DRS-01-05, Project Number 101073924 (ONELAB) for funding Dr L.S. Petralia. A. Chawaguta acknowledges the FFG IKT der Zukunft, IKT der Zukunft, IKT der Zukunft-Resilienz und Distancing (DEVICE) for funding his PhD position.

## Data availability

All data used in this publication are readily available upon request from the corresponding author.

## Ethics declaration

This study was approved by the Institutional Ethics Committee of the Medical University Innsbruck. The study protocol conforms to the ethical guidelines of the 1975 Declaration of Helsinki. Written informed consent was obtained from all participants prior to their inclusion in the study.

