## Supplementary Material for "Sweet interference: oral fermentation volatile confounders in exhaled breath revealed by minimal glucose exposure"

### Targeted VOCs

PCA was performed on a targeted VOCs panel comprising limonene (contained in the administered candies), ethanol, acetic acid, acetoin, and 2,3‑butanediol (glucose‑derived fermentation products), supplemented by acetone and isoprene as ubiquitous, well‑characterized systemic breath markers.

| **Basis for inclusion** | **Compound(s)** | **Source / justification** |
| --- | --- | --- |
| **Untargeted PCA of prior GC-IMS oral/nasal study** | Ethanol  Acetoin  Acetone  Isoprene | Identified as the high-magnitude loading vectors in a genuinely untargeted (hypothesis-free) principal component analysis of paired oral and nasal end-tidal exhaled breath from twenty-one volunteers [14]. Ethanol and acetoin are strongly oral-route-dominated; acetone and isoprene show no significant route dependence and hence serve as systemic reference compounds. |
| **Glycolysis/Fermentation-pathway (domain) knowledge** | Acetic acid  2,3-Butanediol | Added on biochemical grounds as the expected products of two named branches of pyruvate/Embden-Meyerhof-Parnas fermentation are (i) acetic acid from the mixed-acid branch, and (ii) 2,3-butanediol from reduction of acetoin by butanediol dehydrogenase (see main manuscript Table 2). |
| **Study protocol (exogenous marker)** | Limonene | Monitored as an exogenous marker of the lemon-flavoured glucose vehicle, providing an internal indicator of local oral release. Dictated by the intervention protocol rather than selected as a candidate biomarker. |

**Table S1.** The seven target VOCs discussed in the main text (limonene, ethanol, acetic acid, acetoin, 2,3-butanediol, acetone and isoprene) were selected a priori on the basis of their established relevance to oral fermentation chemistry and systemic breath physiology, as set out in the Introduction and in Table 2.

### Compound identification

Compound identities were assigned by comparison with authentic standards. GC‑IMS provides chromatographic and product ion mobility separation The operating conditions used for the preliminary GC-IMS measurements were identical to those reported in our previous study [14]. Figure S1 and Table S2 summarise the assignments of acetoin, acetic acid and limonene, which were confirmed by comparison with the corresponding authentic standards analysed under identical GC IMS conditions. Illustrative zoomed in GC-IMS spectra from three volunteers, obtained before and after the intervention, are presented in Figure S3.

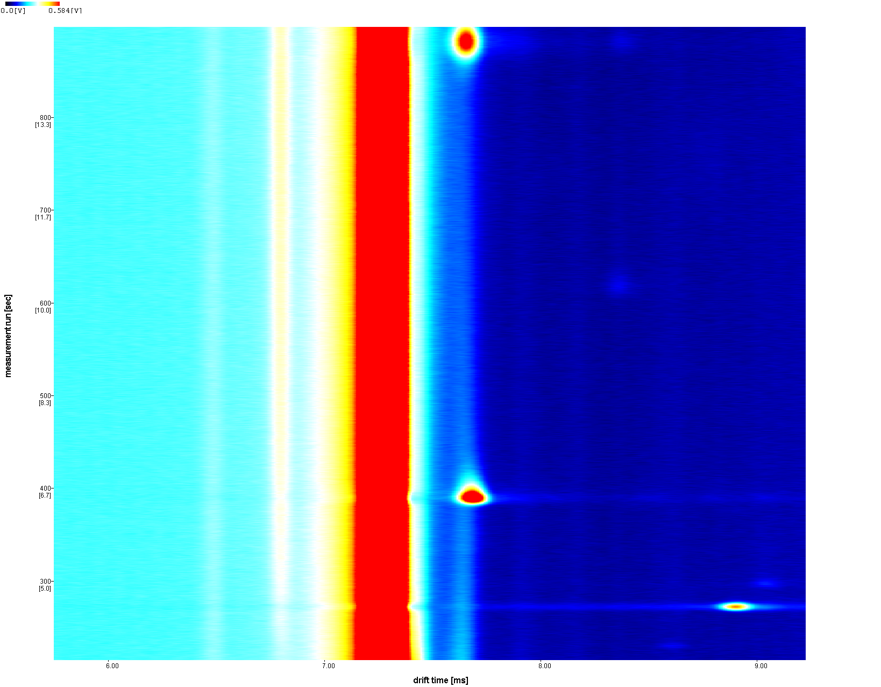

**GC Retention time (s)**

**600 -**

**800 -**

**400 -**

**Drift time (ms)**

**6.00**

**7.00**

**8.00**

**9.00**

**1**

**2**

**3**

**Figure S1**. GC-IMS spectrum measuring an injected standard, showing the peak areas of limonene (1), acetoin (2), and acetic acid (3).

| **Compound** | **Formula** | **PTR-ToF-MS** | | **GC-IMS** | | | **Identification basis / notes** |
| --- | --- | --- | --- | --- | --- | --- | --- |
|  |  | **Protonated ion** | **m/z** | **Retention Time (s)** | **Drift time**  **(ms)** | **K0 values**  **cm^2^V^-1^s^-1^** |  |
| **Limonene** | **C_10_H_16_** | **C_10_H_17_^+^** | 137.134 | 273.77 | 8.898 | 1.6993 | Exogenous marker of the flavoured glucose vehicle. PTR-MS cannot distinguish limonene from other monoterpene isomers. GC-IMS retention time and drift time, matched against measured limonene standard (CAS 5989-27-5). |
| **Acetoin** | **C_4_H_8_O_2_** | **C_4_H_9_O_2_^+^** | 89.060 | 393.82 | 7.682 | 1.9683 | Exactly isobaric with ethyl acetate (constitutional isomers). Discrimination rests on the GC-IMS retention time and drift time, matched against an authentic acetoin standard (CAS 513-86-0). |
| **Acetic acid** | **C_2_H_4_O_2_** | **C_2_H_5_O_2_^+^** | 61.029 | 882.47 | 7.682 | 1.9683 | Discrimination rests on the GC-IMS retention time and drift time, matched against an acetic acid standard (CAS 64-19-7). |

**Table S2** Compound identification table reporting the PTR-ToF-MS and the GC-IMS details: protonated ion m/z, retention time (in seconds), product ion drift times (in ms), and reduced ion mobility K0 (in cm^2^V^-1^s^-1^).

**Pre-intervention**

**Post-intervention**

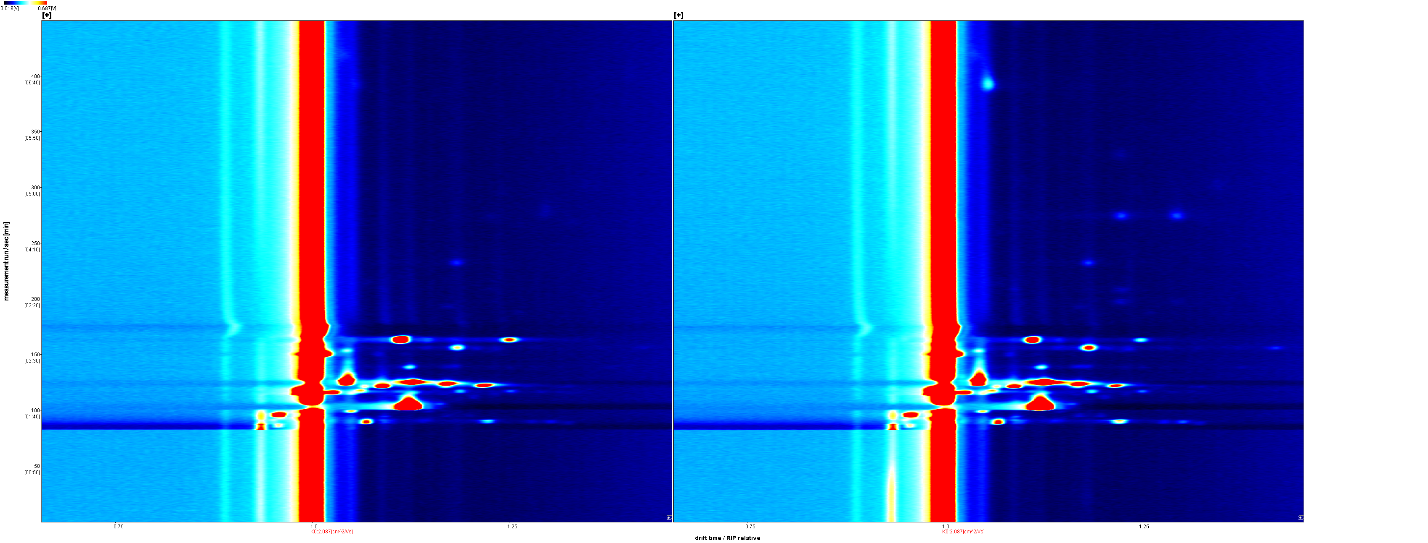

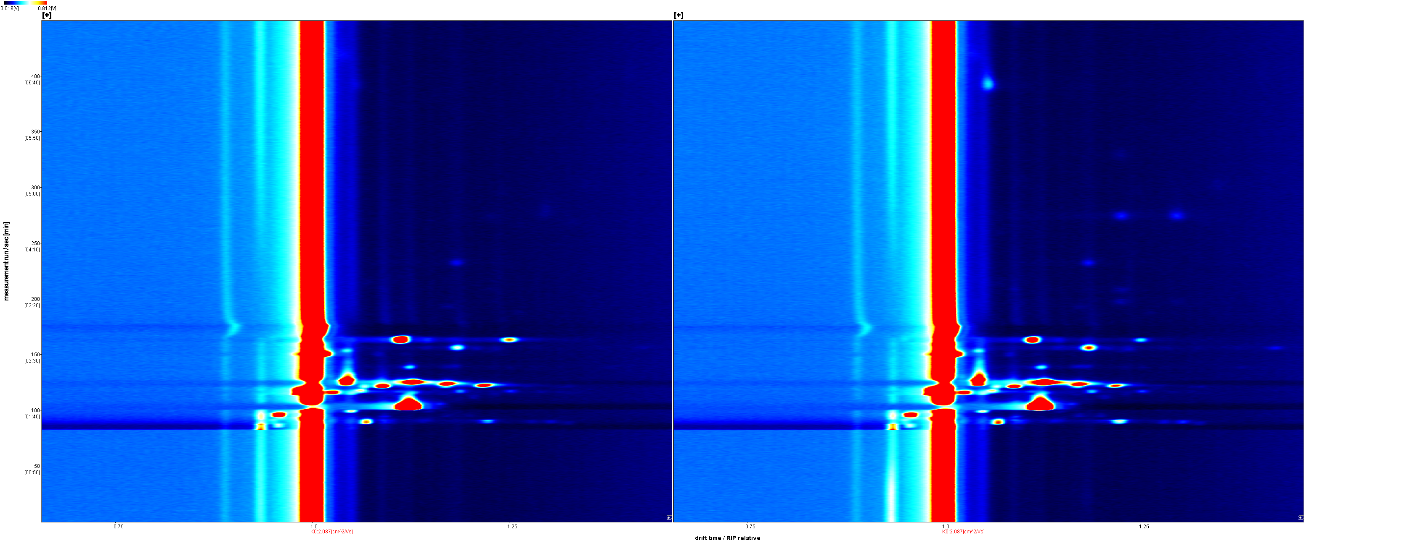

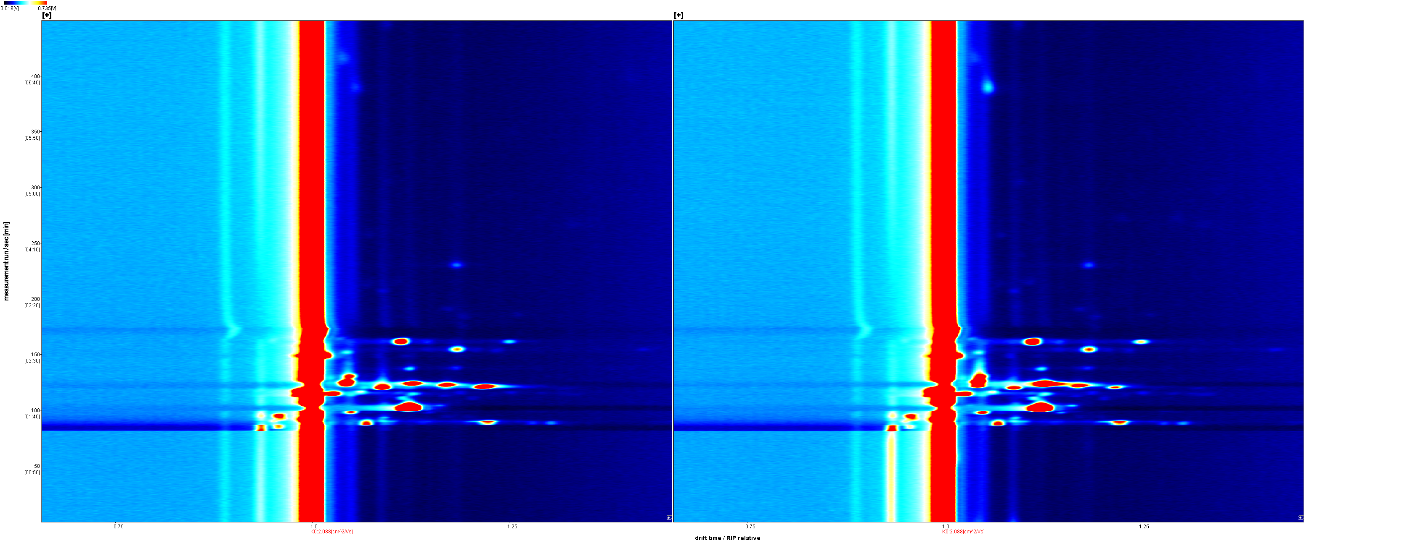

**Volunteer C**

**Volunteer B**

**Volunteer A**

**GC Retention time (s)**

**100 -**

**200 -**

**300 -**

**400 -**

**GC Retention time (s)**

**100 -**

**200 -**

**300 -**

**400 -**

**GC Retention time (s)**

**100 -**

**200 -**

**300 -**

**400 -**

**Drift time RIP-relative**

**0.75**

**1.25**

**1.00**

**0.75**

**1.25**

**1.00**

**Drift time RIP-relative**

**Figure S2** Illustrative GC-IMS zoomed-in spectra for pre- versus post-intervention performed for three volunteers before switching to a study protocol using PTR‑ToF‑MS to achieve high‑temporal‑resolution. In each of the post-intervention panel, the appearing acetoin peak is highlighted with an arrow.
